# A self-amplifying RNA vaccine against COVID-19 with long-term room-temperature stability

**DOI:** 10.1101/2022.03.22.485230

**Authors:** Emily A. Voigt, Alana Gerhardt, Derek Hanson, Peter Battisti, Sierra Reed, Jasneet Singh, Raodoh Mohamath, Madeleine F. Jennewein, Julie Bakken, Samuel Beaver, Christopher Press, Patrick Soon-Shiong, Christopher J. Paddon, Christopher B. Fox, Corey Casper

## Abstract

mRNA vaccines were the first to be authorized for use against SARS-CoV-2 and have since demonstrated high efficacy against serious illness and death. However, limitations in these vaccines have been recognized due to their requirement for cold storage, short durability of protection, and lack of access in low-resource regions. We have developed an easily-manufactured, potent self-amplifying RNA (saRNA) vaccine against SARS-CoV-2 that is stable at room temperature. This saRNA vaccine is formulated with a nanostructured lipid carrier (NLC), providing enhanced stability, improved manufacturability, and protection against degradation. In preclinical studies, this saRNA/NLC vaccine induced strong humoral immunity, as demonstrated by high pseudovirus neutralization titers to the Alpha, Beta, and Delta variants of concern and induction of long-lived bone marrow-resident antibody secreting cells. Robust Th1-biased T-cell responses were also observed after prime or homologous prime-boost in mice. Notably, the saRNA/NLC platform demonstrated thermostability at room temperature for at least 6 months when lyophilized. Taken together, this saRNA delivered by NLC represents a potential improvement in RNA technology that could allow wider access to RNA vaccines for the current COVID-19 and future pandemics.

## INTRODUCTION

More than 470 million confirmed cases of coronavirus disease 2019 (COVID-19) and 6 million related deaths have occurred worldwide due to severe acute respiratory syndrome coronavirus 2 (SARS-CoV-2) as of March 2022.^1^ Although multiple SARS-CoV-2 vaccines have been developed and over 11 billion SARS-CoV-2 vaccine doses have been administered globally,^1^ only 14.4% of people in low-income countries have received a dose.^2^ New vaccines that can be manufactured rapidly and stored without complex cold-chain requirements and result in long-term, robust, multi-variant immunity are urgently needed to protect people across the globe against SARS-CoV-2.

Since the emergence of SARS-COV-2 in 2019, two mRNA vaccines, Moderna mRNA-1273 and Pfizer-BioNTech BNT162b2, were developed most rapidly and proved to be most efficacious^3–5^. These vaccines use nucleoside-modified mRNAs encoding the prefusion-stabilized spike protein of SARS-CoV-2, which are encapsulated by lipid nanoparticles (LNPs) for stabilization and intracellular delivery.^6, 7^ However, global access to these COVID-19 vaccines has been severely hampered by the complex cold-chain requirements for mRNA vaccines and limited vaccine distribution in low- and middle-income countries.^8^ Unopened vials of the currently administered mRNA/LNP vaccines, mRNA-1273 and BNT162b2, are stable for 9 months frozen (-25 to -15°C or -90 to -60°C, respectively), 1 month refrigerated (2-8°C), and 24 or 4 hours at room temperature, respectively.^9, 10^ Despite efforts, long-term stability of mRNA/LNP systems at non-frozen temperatures may be difficult to achieve due to the susceptibility of mRNA to cleavage and the inherent complexity of the LNP delivery system,^11, 12^ and stability is limited even under frozen or dried conditions.^12–14^

Improvements to RNA vaccines are needed to increase stability during storage at temperatures above freezing and to streamline manufacturing processes for global distribution.^15, 16^ Additionally, current mRNA/LNP vaccine technology requires proprietary ionizable lipids, a consistent supply of GMP raw materials for LNP production, and complex manufacturing processes that are difficult to scale for a pandemic response.^17^ Waning of the immune responses to these authorized mRNA/LNP vaccines and the continued emergence of SARS-CoV-2 variants of concern are now necessitating the use of regular booster doses,^18–20^ further hampering full protection of individuals in low-resource countries.^21^ Development of a more potent, easily manufactured, and thermostable RNA vaccine against COVID-19 is necessary to enable global access and a comprehensive pandemic response.

Self-amplifying RNA (saRNA) vaccines—in which RNAs encode viral replicase genes in addition to antigen genes—promise improved potency and durability due to both the amplification of the RNA *in vivo* after delivery and the adjuvanting properties of dsRNA and replication intermediates.^11, 22, 23^ Indeed, saRNAs have been successfully used to create COVID-19 vaccines^24–30^ that have reached clinical trials.^31, 32^ However, thermostability of these vaccine formulations is either unknown^24, 25, 27–30^ or 1 week at refrigerated or room-temperature conditions.^26^ The stability of saRNA vaccines encapsulated in LNPs is unlikely to differ significantly from LNP-encapsulated mRNA vaccines.

We previously developed a lyophilizable, thermostable saRNA vaccine platform with high potency^33^ and long-term storage capability.^34^ This platform is based on a unique, highly manufacturable nanostructured lipid carrier (NLC)^33^ that allows for lyophilization of the drug product, which can extend shelf life to 21 months in refrigerated conditions and 8 months at room temperature.^34^ In the current study, we have applied this thermostable saRNA vaccine platform to COVID-19, efficiently expressing SARS-CoV-2 spike protein antigens and driving induction of robust humoral and cellular SARS-CoV-2- specific immune responses in mice. This formulation is thermostable when lyophilized, maintaining immunogenicity for at least 6 months at room temperature or standard refrigerated conditions. Taken together, this potent and shelf-stable RNA vaccine platform could be an important contribution to increasing global access to COVID-19 vaccines, especially in low-resource regions.

## RESULTS

### saRNA/NLC SARS-CoV-2 vaccine characterization and antigen expression

Our baseline saRNA/NLC SARS-CoV-2 vaccine candidate (designated D614G) consisted of the codon-optimized Wuhan-D614G spike protein sequence^35^ (Figure 1A). This vaccine saRNA was complexed with an NLC delivery nanoparticle by simple mixing to create the vaccine complex with the RNA bound to the exterior of the NLC nanoparticle (Figure 1B) and with an average hydrodynamic particle size of 89 ± 3 nm (Figure 1C). Complexing the saRNA with NLC allowed for retention of RNA integrity even in the presence of RNase (Figure 1D). The formulated vaccine was then transfected into HEK-293 cells where it demonstrated significant SARS-CoV-2 spike protein expression 24 hours post-transfection as measured by western blot (Figure 1E). In order to mitigate supply chain issues for the least readily available ingredient of the NLCs, the cationic lipid DOTAP, we also tested DOTAP from multiple independent manufacturers and demonstrated similar physicochemical properties and 3-month stability of the resulting NLCs, indicating that material to supply large-scale manufacturing may likely be successfully sourced from multiple vendors (Supplementary Table S1).

**Figure 1.**
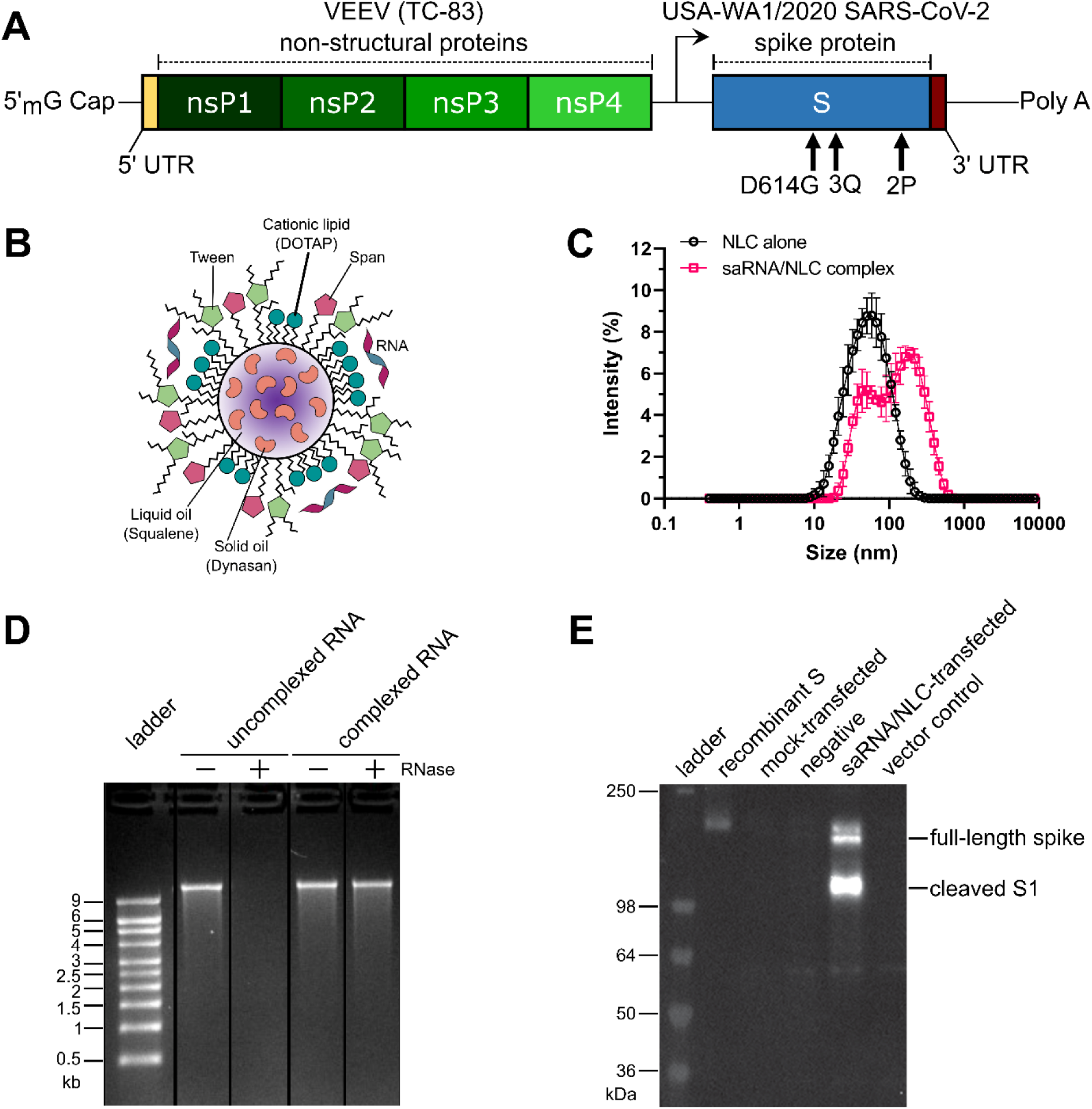
Design and characterization of SARS-CoV-2 D614G saRNA/NLC vaccine. A) SARS-CoV-2 saRNA vaccine schematic. B) RNA/nanostructured lipid carrier (NLC) vaccine formulation. Design by Cassandra Baden. C) Size distribution of NLC formulation particles alone (black) and saRNA/NLC complex (pink). D) SARS-CoV-2 saRNA complexed to the outside of the NLC particles is full-length intact saRNA that is protected by the NLC complexation from RNase degradation. E) Western blot showing SARS-CoV-2 spike (S) protein expression in saRNA/NLC vaccine-transfected HEK-293 cells. See also Figure S1.

### Baseline saRNA/NLC vaccine with the D614G construct induces strong Th1-biased neutralizing antibody responses in mice

To test vaccine immunogenicity of the baseline D614G vaccine candidate, complexed saRNA/NLC was injected intramuscularly into C57BL/6J mice at 1, 10, or 30 µg doses, with a boost dose 3 weeks post-prime. As a negative vector control, an saRNA expressing the non-immunogenic reporter secreted embryonic alkaline phosphatase (SEAP) gene in the place of the SARS-CoV-2 spike gene was used at a 10 µg dose. Serum samples were taken 21 days post-prime and 21 days post-boost. These samples were assessed for SARS-CoV-2 spike protein-binding IgG, IgG1, and IgG2a antibodies by enzyme-linked immunosorbent assay (ELISA) and for SARS-CoV-2 neutralizing antibodies by pseudovirus neutralization assay (Figure 2).

**Figure 2.**
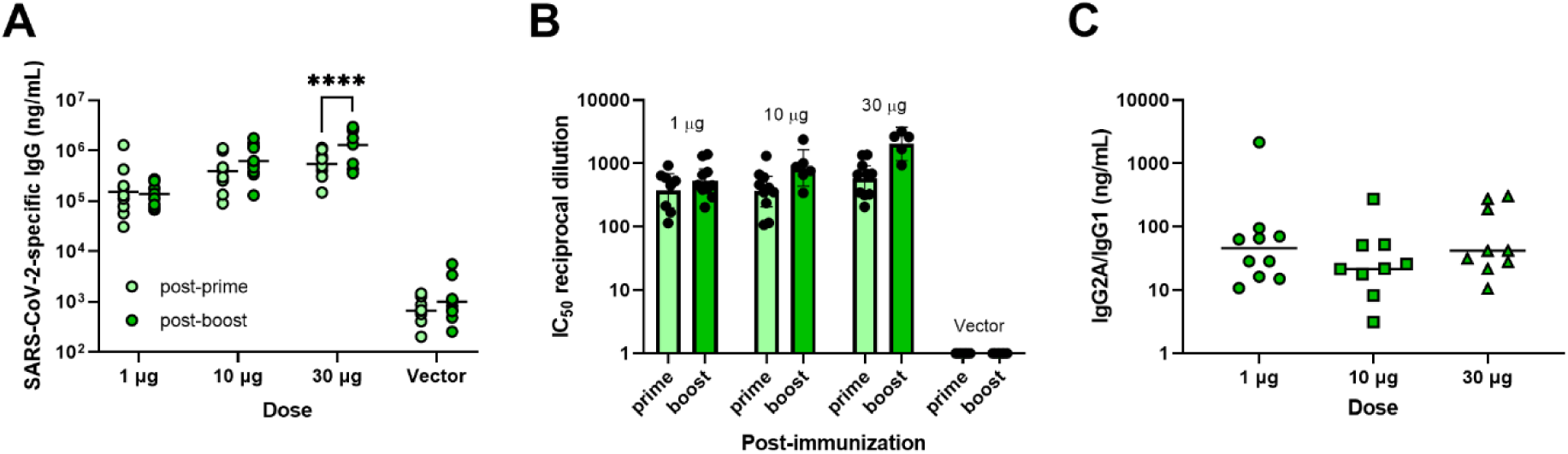
Immunogenicity of SARS-CoV-2 D614G saRNA/NLC vaccine after prime or prime-boost immunization of C57BL/6J mice. A) Serum SARS-CoV-2 spike protein-binding IgG. \*\*\*\**p <* 0.0001. B) Serum SARS-CoV-2 neutralizing antibodies against the original Wuhan strain. C) Serum SARS-CoV-2 spike protein-binding IgG1 versus IgG2a indicates a strong Th1-biased response. The vector control represents mice injected with 10 μg of NLC-complexed saRNA expressing the non-immunogenic secreted embryonic alkaline phosphatase gene.

Strong antibody responses to SARS-CoV-2 spike protein were detected in all vaccinated mice post-prime (Figure 2A), with a statistically significant but relatively small rise in titers after the boost dose (*p* = 0.0054). A clear dose-dependency was seen across the mouse groups. Strong neutralization of Wuhan-D614G pseudotyped lentiviral particles was evident in all vaccinated mouse sera post-prime and post-boost (Figure 2B), with no detectable neutralization noted in vector control-vaccinated mouse sera. Ratios of IgG2a (Th1) to IgG1 (Th2) antibody titers indicated a robust Th1 bias in the humoral responses to the baseline D614G saRNA/NLC vaccine (Figure 2C).

### Antigen optimization increases SARS-CoV-2 variant cross-neutralization

We next sought to optimize the SARS-CoV-2 spike protein sequence for the induction of SARS-CoV-2 antibodies capable of neutralizing multiple variants of the virus. C57BL/6J mice were injected intramuscularly with 10 µg of saRNA/NLC vaccine expressing the native D614G spike protein sequence, the D614G spike protein sequence with a diproline mutation that stabilized the prefusion conformation of the spike trimer^36, 37^ (designated D614G-2P), or the D614G diproline-stabilized spike protein with a QQAQ mutation that ablated the furin cleavage site^38^ (designated D614G-2P-3Q). Each saRNA candidate was complexed with NLC in the same manner as with the baseline candidate, and similar vaccine particle size, RNA integrity, and *in vitro* spike protein expression were observed as in Figure 1 (Supplemental Figure S1). Mice were again administered prime and boost injections 3 weeks apart, and serum samples were taken 3 weeks post-prime and 3 weeks post-boost vaccine dose. All vaccine candidates were well tolerated based on mouse general appearance and behavior, with no signs of weight loss or injection site reactogenicity, similar to previous studies with this saRNA/NLC platform.^33, 34, 39, 40^ SARS-CoV-2 spike-binding IgG antibodies were assessed in mouse sera by ELISA, and SARS-CoV-2 variant neutralization was evaluated by pseudovirus neutralization assay (Figure 3).

**Figure 3.**
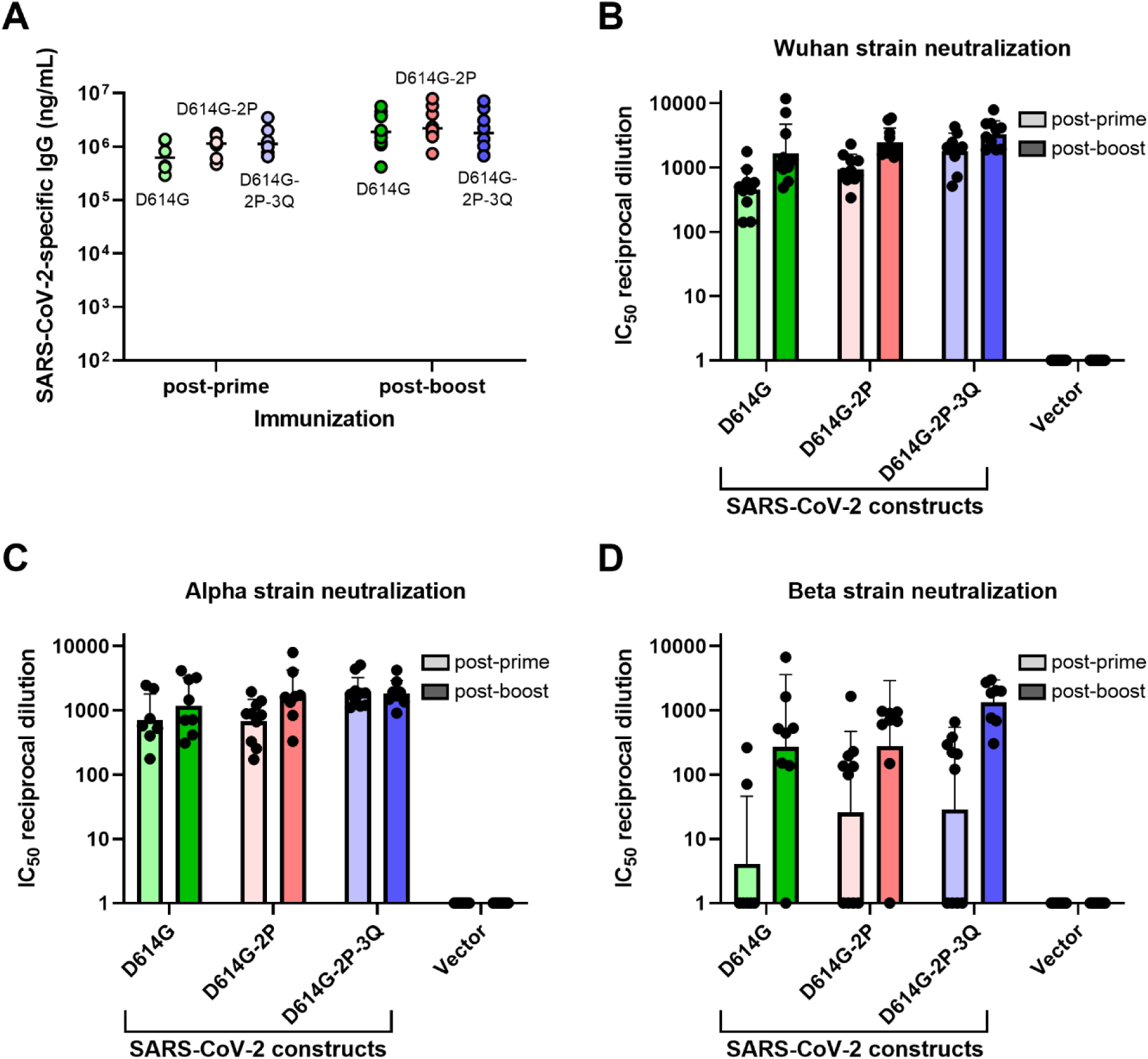
Immunogenicity of SARS-CoV-2 D614G, D614G-2P, and D614G-2P-3Q saRNA/NLC vaccines after 10 µg prime or prime-boost immunization of C57BL/6J mice. A) Serum SARS-CoV-2 spike protein-binding IgG. B-D) Serum SARS-CoV-2 neutralizing antibody titers against the original Wuhan, Alpha (B.1.1.7), and Beta (B.1.351) pseudovirus variants. The vector control represents mice injected with 10 μg of NLC-complexed saRNA expressing the non-immunogenic secreted embryonic alkaline phosphatase gene.

All three vaccine candidates induced strong SARS-CoV-2 spike-binding serum IgG titers at high levels post-prime, with a significant increase post-boost (Figure 3A, average 2.7-fold change post-boost, *p* = 0.013 across groups). No significant difference in spike-binding IgG titers was seen between the three candidates post-boost (*p* = 0.7276). The candidates differed only in their ability to consistently induce antibodies in all mice that cross-neutralize emerging SARS-CoV-2 variants (Figure 3B-D). All three candidates largely retained the neutralization capacity to the Alpha variant (B.1.1.7, UK). However, cross-neutralization against the Beta (B.1.351, South African) variant decreased significantly in D614G- and D614G-2P-vaccinated mouse sera (2.3-fold [*p <* 0.0059] and 4.5-fold [*p =* 0.0010] change post-prime, respectively). This reduction in neutralizing capacity was smaller in sera from D614G-2P-3Q- vaccinated mice and no longer statistically significant (2.2-fold change, *p =* 0.3326), indicating improved induction of variant cross-neutralizing antibodies by this vaccine candidate. The D614G-2P-3Q saRNA/NLC vaccine candidate was therefore chosen as the lead vaccine candidate for further immunogenicity studies and is hereafter referred to as “AAHI-SC2.”

### Optimized saRNA/NLC vaccine AAHI-SC2 induces strong variant cross-protective immune responses and establishment of bone marrow-resident antibody secreting cell populations

After selecting AAHI-SC2 as lead vaccine candidate, we conducted a vaccine dosing study in mice with comprehensive humoral and cellular immunogenicity characterization. C57BL/6J mice were immunized in prime or prime-boost regimens with AAHI-SC2 at doses of 1, 10, or 30 µg. Serum samples were collected to measure SARS-CoV-2 spike-specific IgG (Figure 4A) and neutralizing antibody titers (Figure 4B-C). Bone marrow samples were taken post-boost and used for measurement of SARS-CoV-2 spike-specific IgA- and IgG-secreting cells by ELISpot (Figure 4D-E).

**Figure 4.**
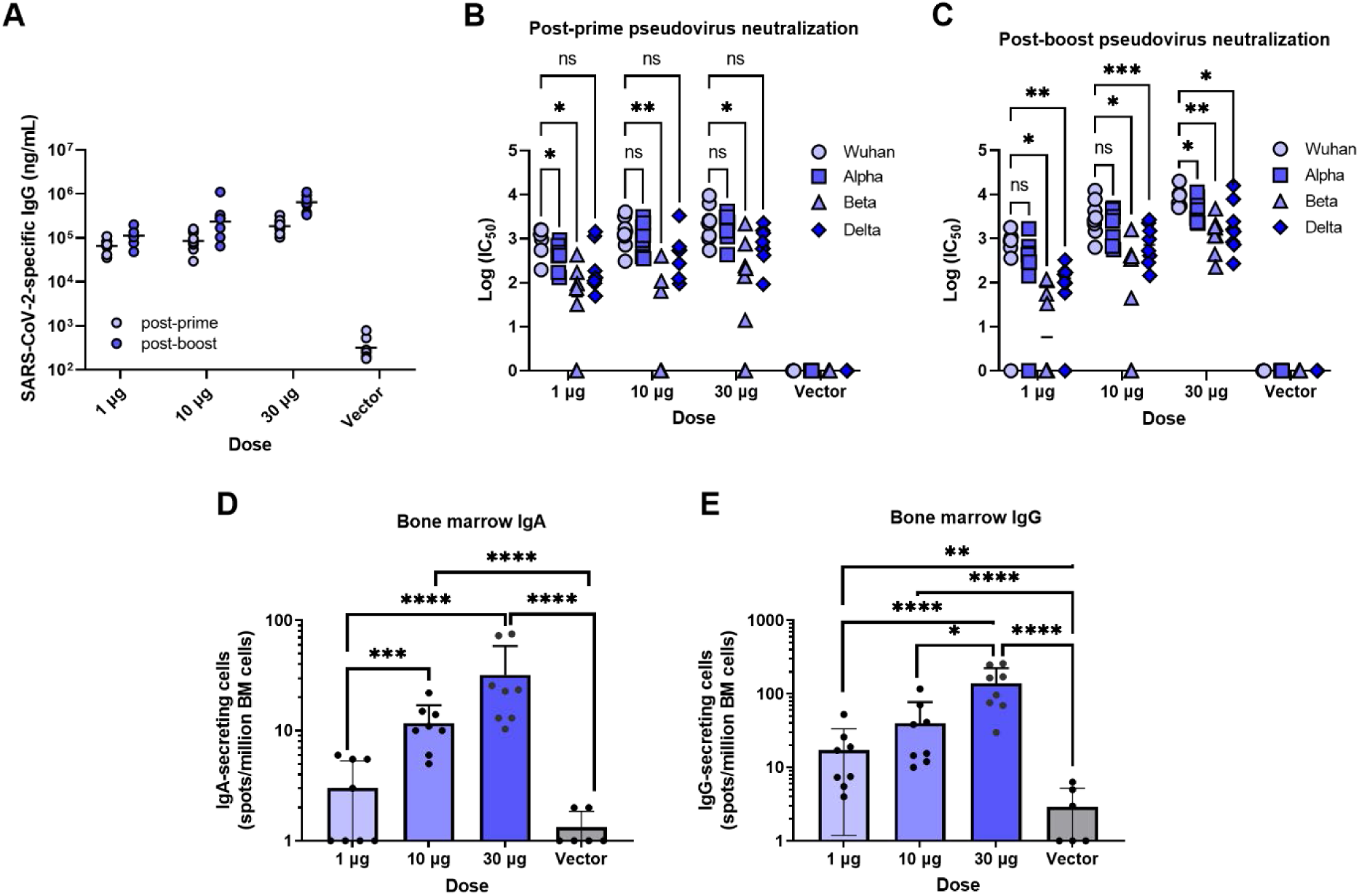
Humoral immunogenicity profiles of optimized SARS-CoV-2 D614G-2P-3Q saRNA/NLC vaccine (AAHI-SC2) after prime or prime-boost immunization of C57BL/6J mice. A) Serum SARS-CoV-2 spike protein-binding IgG. B-C) Serum SARS-CoV-2 neutralizing antibody titers post-prime (B) and post-boost (C). Data were log-transformed and evaluated by mixed effects analysis with multiple comparisons. D-E) Induction of bone marrow (BM)-resident IgA- (D) and IgG-secreting (E) cells by ELISpot. \**p <* 0.05, \*\**p <* 0.01, \*\*\**p <* 0.001, \*\*\*\**p <* 0.0001. The vector control represents mice injected with 10 μg of NLC- complexed saRNA expressing the non-immunogenic secreted embryonic alkaline phosphatase gene.

High levels of SARS-CoV-2 IgG antibodies were induced in mouse sera in a dose-dependent manner, with even a 1 µg dose inducing high responses (Figure 4A). Boost vaccine doses increased IgG titers a small but significant degree at all doses (average 3-fold change, *p <* 0.0001). Sera were able to neutralize Wuhan, Alpha (B.1.1.7), Beta (B.1.351), and Delta (B.1.617.2) pseudoviruses at high levels, both after prime-only and prime-boost vaccination (Figure 4B-C). A significant decrease in the ability of the mouse sera to neutralize the Beta and Delta variants was evident relative to Wuhan strain neutralization as expected both post-prime (average *p <* 0.0001 and *p =* 0.0002, respectively) and post-boost (*p =* 0.0006 and *p <* 0.0001, respectively); however, neutralization of all three variants still generally remained at high, detectable levels. High levels of post-boost bone marrow-resident IgA-secreting and IgG-secreting cells were detected (Figure 4D-E), suggesting induction of at least low levels of mucosal immune responses by the vaccine as well as establishment of robust long-lived IgG-secreting plasma cell populations.

### Optimized AAHI-SC2 vaccine induces robust Th1-biased cellular immune responses

We also characterized cellular responses after prime-only or prime-boost vaccination by stimulation of splenocytes with SARS-CoV-2 overlapping peptides followed by T-cell ELISpot (Figure 5A, B) or intracellular cytokine staining and flow cytometry (Figure 5C-F). Robust Th1-biased T cell responses were seen in vaccinated mice. T cell ELISpots demonstrated large numbers of IFNγ-secreting cells after stimulation with SARS-CoV-2 spike protein overlapping peptides even after a prime dose alone, with clear vaccine dose-dependent responses evident (Figure 5 A). IL-5- (Figure 5B) and IL-17A-secreting (data not shown), Th2-type T cells were induced at very low or undetectable levels, indicating a strong Th1 bias in cellular responses to AAHI-SC2 vaccination. Flow cytometry of splenocytes stimulated with SARS-CoV-2 spike overlapping peptides and stained for a panel of T cell markers and cytokines generated results consistent with the T cell ELISpot findings. High levels of responsive CD4+ Th1 and robust CD8+ T cell populations were stimulated with prime vaccination alone, and these were increased several-fold in a dose-dependent manner after boost injection. Polyfunctional, triple-positive IFNγ-, IL-2-, and TNFα-secreting CD4+ (Figure 5C) and CD8+ T cells (Figure 5D) were stimulated at whole-percentage levels after a prime dose alone and remained high post-boost. Large populations of IFNγ+ and TNFα+ cells post-prime, increasing to a large degree post-boost, drove a significant SARS-CoV-2-specific significant CD4 Th1 response and a large CD8 T cell response (Figure 5E-F), representing up to 20% of the CD8 cell population in the highest-dosed group. Conversely, the CD4 Th2 response was minimal at all timepoints, generally remaining under 2% of CD4 cells, and only measurable at the highest vaccine doses (Figure 5E-F).

**Figure 5.**
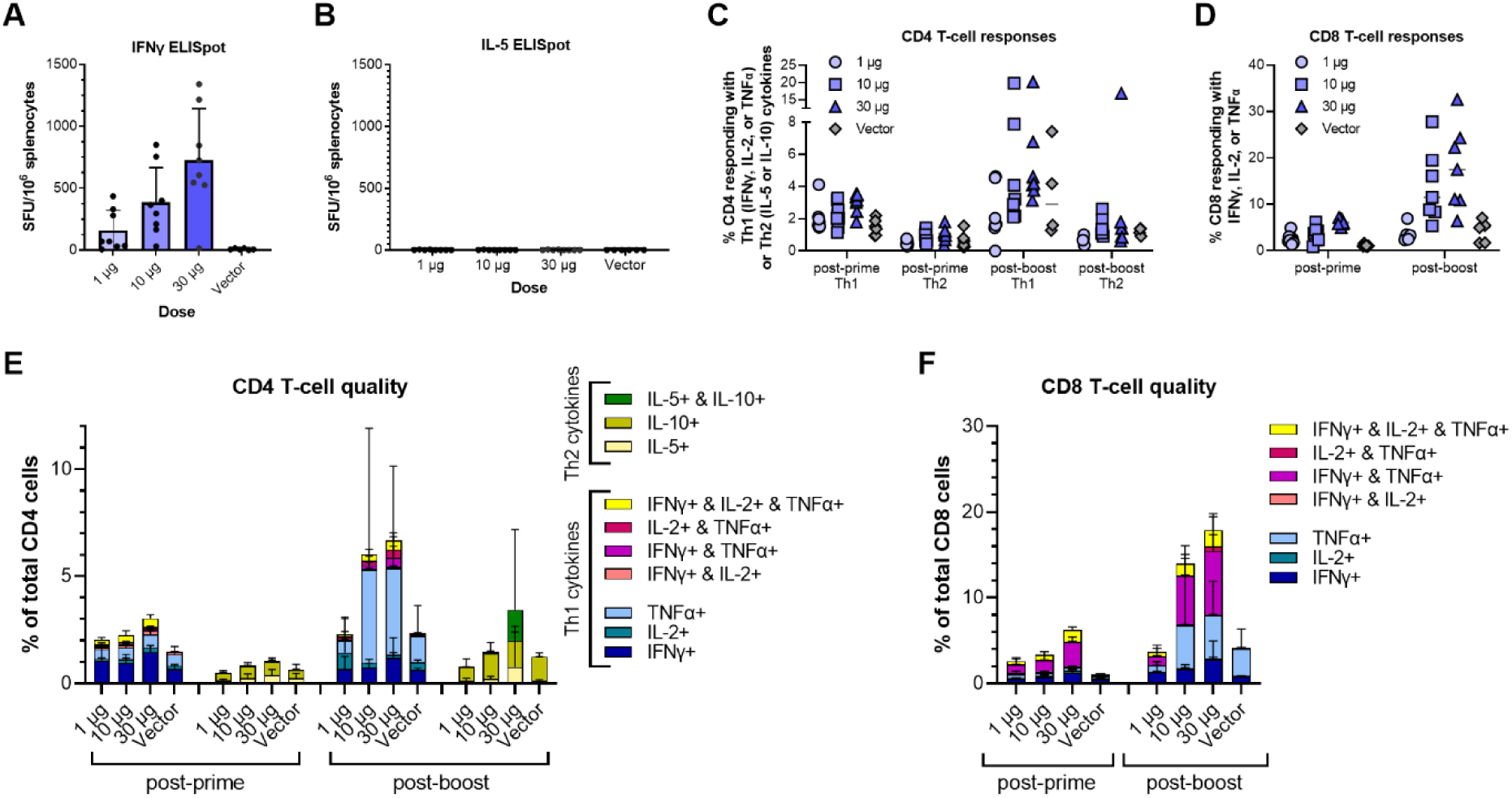
Detailed cellular immunogenicity profiles of optimized SARS-CoV-2 D614G-2P-3Q saRNA/NLC vaccine (AAHI-SC2) after prime or prime-boost immunization of C57BL/6J mice. A) SARS-CoV-2 spike-reactive spleen-resident IFNγ+ T cells by ELISpot. SFU = spot forming units. B) SARS-CoV-2 spike-reactive spleen-resident IL-5+ T cells by ELISpot. C-D) CD4 (C) and CD8 (D) T cells responding with any Th1 (IFNγ, IL-2, or TNFα) or Th2 (IL-5 or IL-10) cytokines post-prime and post-boost were plotted representing the total number of responding cells, for CD4 or CD8 cells per mouse. E-F) Quality of responding CD4 and CD8 T cells. Total Th1 or Th2 responses were subdivided by cells responding with one, two, or three cytokine(s) to show magnitude and quality of Th1 and Th2 responses, with bar graphs showing the average percentage across each group of mice. The vector control represents mice injected with 10 μg of NLC-complexed saRNA expressing the non-immunogenic secreted embryonic alkaline phosphatase gene.

### Lyophilized AAHI-SC2 vaccine is thermostable, enabling long-term storage

Short-term stability of the liquid AAHI-SC2 vaccine in an excipient background of 20% w/v sucrose and 5mM sodium citrate was evaluated over 2 weeks of storage at -80°C, -20°C, 2-8°C, 25°C, and 40°C and compared to freshly prepared controls. Vaccine formulation particle size (70-100 nm) was maintained over the 2-week period at all storage temperatures (Supplemental Figure S2A). The vaccine also retained its ability to induce SARS-CoV-2-specific serum IgG in C57BL/6 mice as measured by ELISA after storage at all temperatures except 40°C (Supplemental Figure S2B) and showed a trend towards potentially reduced potency developing in the 25°C-stored vaccine after 2 weeks.

In order to achieve long-term storage stability, we lyophilized the AAHI-SC2 vaccine and characterized it before and after storage at various temperatures compared to the frozen liquid vaccine. Previously, we demonstrated proof of concept for the long-term storage stability of this platform using SEAP reporter saRNA in the presence of 20% w/v sucrose as a lyoprotectant with a background of 2mM sodium citrate.^34^ Here, the AAHI-SC2 vaccine was lyophilized under those same conditions in addition to two alternative excipient backgrounds: 20% sucrose with 5mM sodium citrate (increasing the concentration of sodium citrate, a metal chelator) and 20% sucrose with 1mM sodium citrate and 10mM Tris (increasing the pH of the system with addition of Tris buffer). Data with the alternative excipient backgrounds are presented in Supplemental Figure S3; however, the original proof-of-concept excipient formulation performed the best (Figure 6).

**Figure 6.**
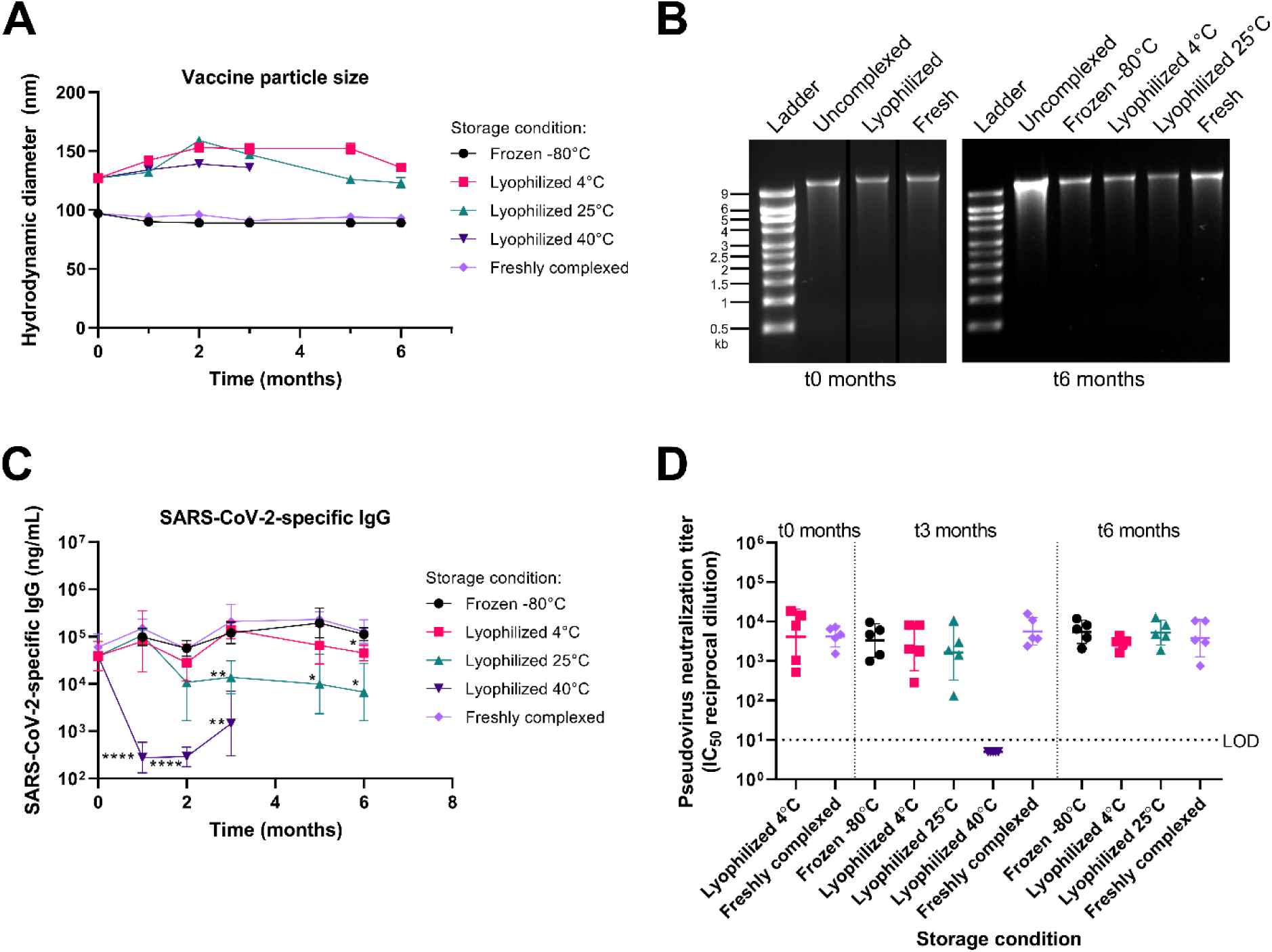
Stability of the lyophilized SARS-CoV-2 saRNA/NLC vaccine (AAHI-SC2) after 6 months of storage at different temperatures. A) Hydrodynamic (Z-average) diameter of the vaccine complex. B) Integrity of vaccine RNA at 0 and 6 months of storage at the indicated temperatures. C) Serum SARS-CoV-2 spike protein-binding IgG induced in female C57BL/6J mice by the vaccine after storage under the indicated conditions for the indicated times. *n* = 5 mice per condition per timepoint. Statistical analysis conducted on log-transformed data by one-way ANOVA at each timepoint, comparing each stored vaccine preparation to freshly complexed vaccine with multiple comparison corrections. \**p <* 0.05, \*\**p <* 0.01, \*\*\*\**p <* 0.0001. D) Pseudovirus neutralization titers induced in female C57BL/6J mice by the vaccine after storage under the indicated conditions for the indicated times. LOD = limit of detection. *n* = 5 mice per condition per timepoint. See also Figure S3.

Vaccine at each excipient condition was stored frozen at -80°C or lyophilized at 4°C, 25°C, or 40°C. A freshly complexed positive control was also prepared at each timepoint for comparison. For the proof-of-concept excipient formulation, vaccine particle size increased by approximately 30 nm immediately after lyophilization, but after that initial increase, particle size was maintained over the 6 months of the study (Figure 6A). RNA integrity was also maintained after lyophilization and throughout 6 months of storage under frozen conditions and lyophilized conditions stored at 4°C and 25°C (Figure 6B). The accelerated storage condition of 40°C was eliminated from the stability study after 3 months due to complete degradation of the RNA (data not shown).

Vaccine immunogenicity during storage was assessed by injecting stored vaccine intramuscularly into 6-8 week old C57BL/6J mice (5 mice per storage condition per timepoint), collecting mouse serum samples 14 days post-immunization, and testing these sera for SARS-CoV-2 spike protein-binding antibodies by ELISA, as described above. The ability of each stored vaccine to induce an immune response *in vivo* was predicted well by the presence of a major intact RNA band by agarose gel electrophoresis. As expected, the -80°C-frozen vaccine maintained the ability to induce SARS-CoV-2-specific IgG throughout the 6-month stability study (Figure 6C) along with equivalent pseudovirus neutralization titers to freshly complexed vaccine at each timepoint (*p* > 0.05) (Figure 6D). After storage at 40°C for 3 months, both antibody titers and neutralizing titers were diminished with lyophilized vaccine (Figure 6C-D), as expected for this accelerated degradation condition.

The lyophilized vaccine stored at 4°C or at 25°C continued to elicit robust serum SARS-CoV-2-specific-IgG titers after 6 months of storage (Figure 6C). The lyophilized 4°C-stored vaccine maintained its ability to stimulate mouse serum IgG over the course of the study, comparable to the lyophilized vaccine prior to storage; although after 6 months the SARS-CoV-2-specific IgG titer induced by the 4°C-stored vaccine was lower than that induced by the freshly complexed control at that timepoint (*p* < 0.05). The 25°C-stored vaccine after 3 months of storage induced a decreased mouse serum IgG titer relative to both the lyophilized vaccine prior to storage and the freshly complexed vaccine at each timepoint (*p* < 0.05). Despite this apparent decrease in total spike-binding IgG, lyophilized vaccine induced equivalent serum pseudovirus neutralization titers relative to fresh-mixed vaccine or lyophilized vaccine prior to storage, with no significant differences observed between neutralizing antibody titers induced by 4°C- and 25°C-stored lyophilized samples relative to those induced by freshly mixed vaccine at each timepoint or relative to those induced by the lyophilized sample prior to storage (*p* > 0.05) (Figure 6D). This suggests the long-term retention of lyophilized vaccine immunogenicity after storage at refrigerated or even ambient temperatures.

## DISCUSSION

AAHI-SC2, combining the potency and self-adjuvanting nature of saRNA with the improved stability and immune-stimulating properties of the NLC formulation, exhibits many characteristics of a “next-generation” SARS-CoV-2 RNA vaccine. Immunization of mice with AAHI-SC2 induces cross-variant neutralizing antibodies, establishes key long-lived bone marrow-resident antibody secreting cell populations, and generates robust Th1-biased polyfunctional T cell responses. AAHI-SC2 exhibits enhanced thermostability, allowing it to be stored at room temperature or refrigerated conditions for at least 6 months—and possibly far longer—while still maintaining the ability to stimulate equivalent neutralizing antibody titers. This vaccine, soon to begin human testing in accelerated Phase I/II clinical trials, demonstrates excellent preclinical immunogenicity, with far simpler manufacturing and cold-chain requirements than mRNA vaccines currently in use, enabling greater global vaccine access and distribution.

The two mRNA vaccines in widespread use rely on LNP-encapsulated nucleoside-modified RNA, and both are thought to be effective via induction of neutralizing antibody titers.^41, 42^ Our vaccine elicited neutralizing antibody responses in mice that well exceeded levels that have been predicted to be protective.^43^ Direct comparisons of our saRNA/NLC vaccine to the approved mRNA vaccines were not possible in this study; however, literature comparisons suggest that, acknowledging minor differences in the immunogenicity assays used, AAHI-SC2 humoral immunogenicity in C57BL/6J mice likely compares favorably with these mRNA vaccines.^6, 7^

While some studies indicate that neutralizing antibody titers may be considered correlates of protection,^44–48^ T cell responses have been shown to be critical for the control and clearance of infection, as well as for mediating protection against emerging variants and preserving long-term immunity.^48–51^ T cell responses appear earlier than neutralizing antibodies following vaccination,^52^ and the timing of their appearance correlates with the attainment of vaccine protective efficacy.^3^ Although emerging variants of SARS-CoV-2 appear to be able to evade vaccine-induced humoral immune responses,^53, 54^ T cell immunity appears to be more durable in response to these variants^55, 56^ likely because T cell epitopes are more highly conserved in emergent SARS-CoV-2 variants than the corresponding B cell epitopes.^56^ Combined, the literature presents a strong case for the importance of vaccines that induce not only high levels of neutralizing antibodies but also strong and enduring T cell responses. The Moderna and Pfizer-BioNTech vaccines elicit strong levels of neutralizing antibodies,^6, 57^ but the T cell responses to these existing vaccines are less than robust in both preclinical and clinical studies.^6, 57^ AAHI-SC2 appears to elicit strong T cell responses in preclinical studies, possibly due to the combination of the adjuvanting properties of squalene and saRNA, and may provide a foundation for durable and long-term T-cell immunity. Upcoming studies in non-human primates and clinical trials in humans should provide additional data to evaluate the immunogenicity, and particularly the T cell induction, of AAHI-SC2 for comparison to published data on mRNA-1273, BNT162b2, and other SARS-CoV-2 mRNA^6, 7, 58^ or saRNA^24, 25, 27, 29, 30^ vaccines in development.

Th1-biased vaccine immune responses are key for both durable protection and vaccine safety.^59^ Th2 responses have been associated with immunopathology induced by vaccines^41, 59, 60^ or by natural infection with SARS-CoV-2,^61^ while Th1-type responses are associated with durable vaccine immunity^62^ and a lack of observed immunopathology in COVID-19 patients.^62, 63^ It is unclear how widely Th2-type vaccine responses induced by coronavirus vaccines could lead to clinically meaningful immune complications, with no evidence of vaccine-associated enhanced respiratory disease (VAERD) or antibody-dependent enhancement (ADE) of virus infection and disease reported in any human trials for SARS-CoV-1, SARS-CoV-2, or Middle East Respiratory Syndrome (MERS) vaccines.^64, 65^ Regardless, the AAHI-SC2 vaccine clearly induces strong Th1-biased immunity suggesting a favorable immunogenicity profile for a SARS-CoV-2 vaccine.

In addition to its immunogenicity, thermostability of the AAHI-SC2 vaccine is critical for its potential utility. Several components of mRNA vaccines may affect their stability during storage: LNP formulation, modified nucleosides, GC content, RNA length, and non-coding regions.^24, 25^ While optimization of these properties has led to improved mRNA stability, stringent cold-chain requirements are still necessary for the Moderna and Pfizer-BioNTech vaccines.^9, 10^ These requirements were a monumental challenge in the early days of vaccine rollout and continue to be a major impediment to worldwide distribution,^66^ particularly given the short stability window of the Moderna and Pfizer-BioNTech vaccines at room temperature (20-25°C).

Additional RNA vaccines are currently under development with reported stability data, but these data are sparse and indicate limited, short-term room-temperature stability of days to a few weeks at best.^40, 58, 67^ As these reports illustrate, maintenance of RNA integrity during storage and after administration is challenging. RNA molecules are more susceptible to degradation than DNA due to the presence of the 2’ hydroxyl group on the ribose backbone. RNAs must also be protected from degradation by ubiquitous ribonucleases. Lyophilization is a known strategy to stabilize RNA by itself for long-term storage;^68^ however, limited stability has been demonstrated for lyophilized RNA vaccines encapsulated in LNPs.^12, 13^ The complex structure of LNPs may make lyophilization difficult due to the potential for rupture of the lipid layers and/or payload leakage, particularly important due to the encapsulation of the RNA. In contrast, the NLC delivery vehicle, which has a structure more similar to an oil-in-water emulsion than an LNP, has previously demonstrated stability upon lyophilization.^34^ In this work, demonstrating at least 6 months of stability at ambient temperature, the AAHI-SC2 vaccine represents a potential major improvement in RNA vaccine stability and distribution.

Besides its thermostability profile, AAHI-SC2 also offers several manufacturing and supply-chain advantages relative to existing vaccines and vaccine candidates. NLC manufacturing relies on equipment already utilized in standard oil-in-water emulsion-based vaccine adjuvant production and materials that can be sourced from multiple different commercial vendors. In contrast to LNPs, which require encapsulation of the RNA during production, our vaccine technology allows for the NLC and RNA to be produced separately and complexed by simple mixing at any point up to just before administration. This approach greatly simplifies manufacturing, enables stockpiling of NLC for pandemic preparedness,^34^ and allows for post-manufacturing substitution of saRNA species to address emerging viruses or viral variants of concern. The AAHI-SC2 vaccine also does not contain pseudouridine components or highly specialized proprietary lipids whose scarcity and proprietary nature present challenges for sourcing.^17^

## Summary

We present a highly potent SARS-CoV-2 saRNA vaccine that induces strong immune responses in mice and demonstrates establishment of long-lived antibody-secreting plasma cell populations. The potency of this vaccine in conjunction with its induction of broadly cross-variant neutralizing antibodies may allow this vaccine to provide effective protection against many SARS-CoV-2 strains. While this vaccine candidate is only now advancing into non-human primate studies, animal efficacy testing, and human trials, we believe it represents an important advance in RNA vaccine technology, particularly for rural areas and the developing world. A key advantage of this technology is its improved manufacturability and thermostability relative to other mRNA vaccines, enabling lower costs and widespread distribution in lower income countries without the need for an extreme cold chain. This vaccine is scheduled to advance into efficacy studies in non-human primates and an accelerated Phase I/II clinical trial. This thermostable saRNA/NLC vaccine technology is valuable not just for the current SARS-CoV-2 pandemic but also for future pandemic preparedness and responsiveness.

## MATERIALS AND METHODS

### saRNA expression plasmid design, cloning, and production

Three saRNA plasmids each with a unique candidate SARS-CoV-2 spike open reading frame were created, along with a fourth saRNA plasmid expressing secreted embryonic alkaline phosphatase (SEAP) instead of a vaccine antigen as an appropriate vector control. The SARS-CoV-2 spike open reading frame sequence from GenBank MT246667.1 incorporating the additional D614G mutation was used as the “baseline” sequence, with additional modifications to create two additional vaccine candidates: D614G-2P represents the baseline sequence with a substitution of PP for KV at amino acid positions 987-988 and an addition of nine N-terminal codons from the reference genome encoding amino acid sequence MFLLTTKRT; D614G-2P-3Q refers to the baseline sequence with the diproline substitution and an additional substitution of QQAQ for RRAR at the furin cleavage site at amino acid positions 683-686. These sequences were then codon-optimized for mammalian (human) expression by Codex DNA (San Diego, CA) using a proprietary algorithm, synthesized by BioXp (Codex DNA), and inserted into AAHI’s backbone saRNA expression vector by Gibson cloning. The SEAP-expressing plasmid was created by a similar process to insert the SEAP expression sequence in place of the vaccine antigen. Plasmid sequences were all confirmed by Sanger sequencing. Template plasmids were amplified in *E. coli* and extracted using Qiagen (Germantown, MD) maxi- or gigaprep kits, followed by *Not*I linearization (New England Biolabs [NEB], Ipswich, MA). Linearized DNA was purified by Qiagen DNA purification kits.

### RNA manufacture

Vaccine saRNA was generated by T7 polymerase-mediated *in vitro* transcription (IVT) using *Not*I-linearized DNA plasmids as templates. An in-house optimized IVT protocol was used with commercially available rNTP mix (NEB) and commercially available T7 polymerase, RNase inhibitor, and pyrophosphatase enzymes (Aldevron, Fargo, ND). DNA plasmids were digested away (DNase I, Aldevron), and Cap 0 structures were added to the transcripts by treatment with guanylyltransferase (Aldevron), GTP, and S-adenosylmethionine (NEB). RNA was chromatographically purified using Capto Core 700 resin (GE Healthcare, Chicago, IL) followed by diafiltration and concentration through tangential flow filtration using a 750 kDa molecular weight cut-off (MWCO) modified polyethersulfone (mPES) membrane (Repligen, Waltham, MA). Terminal filtration of the saRNA material was done using a 0.22 µm PES filter, and the saRNA materials were stored at −80°C until use/complexation. Agarose gel electrophoresis was used to characterize saRNA size and integrity, and RNA concentration was quantified by UV absorbance (NanoDrop 1000) and RiboGreen assay (Thermo Fisher Scientific, Waltham, MA).

### NLC manufacture

Nanostructured lipid carrier (NLC) formulation was produced as described previously.^33, 34, 39^ Briefly, trimyristin (Sigma-Aldrich, St. Louis, MO), squalene (SAFC Supply Solutions, St. Louis, MO), sorbitan monostearate (Spectrum Chemical Mfg. Corp., New Brunswick, NJ), and the cationic lipid DOTAP (Lipoid, Ludwigshafen, Germany) were mixed and heated at 60°C in a bath sonicator to create the oil phase. Polysorbate 80 (MilliporeSigma, Burlington, MA) was separately diluted in 10mM sodium citrate trihydrate and heated to 60°C in a bath sonicator to create the aqueous phase. After dispersion of components in each phase, a high-shear mixer (Silverson Machines, East Longmeadow, MA) was used at ∼5,000 rpm to mix the oil and aqueous phases. Particle size of the mixture was then further decreased by high-pressure homogenization by processing at 30,000 psi for ten discrete passes using an M110P Microfluidizer (Microfluidics, Westwood, MA). The NLC product was then filtered through a 0.22 µm PES filter and stored at 2°C–8°C until use.

### Vaccine complexing and characterization

Vaccine complexes for immediate *in vivo* injection were created by mixing aqueous RNA 1:1 by volume with NLC diluted in a buffer containing 10mM sodium citrate and 20% w/v sucrose. All vaccines were prepared at a nitrogen:phosphate (N:P) ratio of 15, representing the ratio of amine groups on the NLC DOTAP to phosphate groups on the RNA backbone. This produced a vaccine solution containing the intended dose of complexed saRNA/NLC in an isotonic 10% w/v sucrose, 5mM sodium citrate solution. Vaccine was incubated on ice for 30 minutes after mixing to ensure complete complexing.

Hydrodynamic diameter (particle size) of the NLC and vaccine nanoparticles was determined using dynamic light scattering (Zetasizer Nano ZS, Malvern Panalytical, Malvern, UK) on triplicate 1:100 dilutions in nuclease-free water. Sizing was done in a disposable polystyrene cuvette using previously established parameters.^34^

Free and complexed RNA integrity and NLC-provided protection against RNases were evaluated by visualizing RNA integrity after agarose gel electrophoresis. RNA samples were diluted to 40 ng/μL RNA in nuclease-free water. For RNase treatment, this diluted RNA was incubated with RNase A (10 mg/mL, Thermo Scientific, Waltham, MA) at a mass ratio of 1:500 RNase:RNA for 30 minutes at room temperature, followed by incubation with proteinase K (∼20 mg/mL, Thermo Scientific) at a mass ratio of 1:100 RNase A:proteinase K for 10 minutes at 55°C. For complexed samples, RNA was extracted from complexes prior to gel electrophoresis by phenol:chloroform extraction. All RNA samples were mixed with glyoxal loading dye (Invitrogen, Waltham, MA) 1:1 by volume, incubated at 50°C for 20 minutes, loaded on a denatured 1% agarose gel in NorthernMax-Gly running buffer (Invitrogen) alongside Millennium RNA Markers (Thermo Fisher Scientific), and run at 120 V for 45 minutes before imaging on a ChemiDoc MP Imaging System (Bio-Rad Laboratories, Hercules, CA).

### Vaccine lyophilization

Vaccine complexes intended for lyophilization and storage were prepared similarly to above by mixing aqueous RNA 1:1 by volume with NLC diluted in either 20% w/v sucrose only, 20% w/v sucrose and 5mM sodium citrate, or 20% w/v sucrose with both 1mM sodium citrate and 10mM Tris. All vaccines with all excipient conditions were prepared at an N:P ratio of 15 and were incubated on ice for 30 minutes after mixing to ensure complete complexing. After complexing, vaccine was lyophilized using a VirTis AdVantage 2.0 EL-85 (SP Industries, Warminster, PA) benchtop freeze dryer controlled by the microprocessor-based Wizard 2.0 software. The lyophilization cycle began with a freezing step at −50°C, followed by primary drying at −30°C and 50 mTorr, and finishing with secondary drying at 25°C and 50 mTorr. When the cycle was complete, the samples were brought to atmospheric pressure, blanketed with high-purity nitrogen, and stoppered before being removed from the freeze-dryer chamber. At each timepoint, lyophilized samples were reconstituted using nuclease-free water to their original concentration.

### Mouse studies

All animal work was done under the oversight of the Bloodworks Northwest Research Institute’s Institutional Animal Care and Use Committee (Seattle, WA). All animal work was in compliance with all applicable sections of the Final Rules of the Animal Welfare Act regulations (9 CFR Parts 1, 2, and 3) and the *Guide for the Care and Use of Laboratory Animals, Eighth Edition*.^69^

C57BL/6J mice obtained from The Jackson Laboratory (Harbor, ME) were used for all animal studies in this work. Mice were between 6 and 8 weeks of age at study onset. Mice were immunized by intramuscular injection bilaterally in the rear quadriceps muscle (50 µL/leg, 100 µg total). Survival serum samples were taken by retro-orbital bleed.

### Serum IgG, IgG1, and IgG2a titers by ELISA

SARS CoV-2 spike protein-binding IgG antibodies in mouse serum were measured by ELISA. Plates (384- well high-binding plates, Corning, Corning, NY) were coated with 1 µg/mL of Recombinant SARS-CoV-2 Spike His Protein, Carrier Free, (R&D Systems, Minneapolis, MN; #10549-CV) in phosphate-buffered saline (PBS) and incubated overnight at 4°C. The coating solution was removed, and blocking buffer (2% dry milk, 0.05% Tween 20, and 1% goat serum) was applied for at least 1 hour. On a separate, low-binding plate, each sample was diluted 1:40 and then serially 1:2 to create a 14-point dilution curve for each sample. Naïve mouse serum, used as the negative control, was diluted identically to the samples. A SARS-CoV-2 neutralizing monoclonal antibody (mAb; GenScript, Piscataway, NJ; #A02057), was used as a positive control at a known starting concentration of 3.2 ng/µL followed by serial 1:2 dilutions similarly to each sample and negative control. SARS-CoV-2 spike protein-coated and blocked assay plates were washed, and serially diluted samples were then transferred onto the coated plates followed by a 1-hour incubation. Plates were then washed, and spike protein-bound antibodies were detected using an Anti-Mouse IgG (Fc Specific)-Alkaline Phosphatase antibody (Sigma-Aldrich, #A2429) at a 1:4000 dilution in blocking buffer. Plates were washed and then developed using phosphatase substrate (Sigma-Aldrich, #S0942) tablets dissolved in diethanolamine substrate buffer (Fisher Scientific, Waltham, MA; #PI34064) at a final concentration of 1 mg/mL. After a 30-minute development, plates were read spectrophotometrically at 405 nm. A 4-point logistic curve (4PL) was used to fit the antibody standard titration curve. Sample concentrations were interpolated off the linear region of each sample dilution curve using the standard curve for absolute quantification of antibody titers. For Supplemental Figure S2 only, antibody level was quantified by endpoint titer, with the endpoint titer defined as the dilution at which each sample dilution curve rises to above 3 standard deviations above assay background.

For IgG1 and IgG2a isotype-specific ELISAs, the identical plate coating and blocking procedures were conducted. For the IgG1 assay, the standard curve was run using an IgG1 SARS-CoV-2 neutralizing antibody (GenScript) for full quantification. Sample dilution and incubation were identical to the total IgG curve, and plates were probed with IgG1- and IgG2a-specific secondary alkaline phosphatase-conjugated detection antibodies prior to development, reading, and quantification as described above.

### Pseudovirus neutralization assay

The SARS-CoV-2 pseudovirus neutralizing antibody titers of immunized mouse sera were measured by pseudovirus neutralization assays using procedures adapted from Crawford et al.^70^ Lentiviral SARS-CoV-2 spike protein pseudotyped particles were prepared by following the Bioland Scientific (Paramount, CA) BioT plasmid transfection protocol. Briefly, HEK-293 cells (American Type Culture Collection, Manassas, VA; #CRL-3216) were plated at 4 x 10^5^ cells/mL in 6-well plates 18-24 hours before the assay to achieve 50-70% confluency at assay start. Cellular growth medium was then replaced with serum-free Gibco Dulbecco’s Modified Eagle Medium (DMEM) with GlutaMAX immediately prior to transfection. The HEK-293 cells were co-transfected with several plasmids: a plasmid containing a lentiviral backbone expressing luciferase and ZsGreen (BEI Resources, Manassas, VA; #NR-52516), plasmids containing lentiviral helper genes (BEI Resources; #NR-52517, NR-52518, and NR-52519), and a plasmid expressing a delta19 cytoplasmic tail-truncated SARS-CoV-2 spike (Wuhan strain, B.1.1.7, and B.1.351 plasmids from Jesse Bloom, Fred Hutchinson Cancer Research Center, Seattle, WA; and B.1.617.2 plasmid from Thomas Peacock, Imperial College London, UK). BioT transfection reagent (Bioland Scientific) was used to mediate the co-transfection. The assay plates were incubated for 24 hours at standard cell culture conditions (37°C and 5% CO_2_). Then the serum-free media was aspirated off, and fresh growth medium (Gibco DMEM with GlutaMAX and 10% fetal bovine serum [FBS]) was added to the plates before returning them to the incubator for an additional 48 hours. Pseudovirus stocks were collected by harvesting the supernatants from the plates, filtering them through a 0.2 μm PES filter (Thermo Scientific), and freezing at -80°C until titration and use.

Mouse serum samples were diluted 1:10 in medium (Gibco DMEM with GlutaMAX and 10% FBS) and then serially diluted 1:2 for 11 total dilutions in 96-well V-bottom plates. Polybrene (Sigma-Aldrich) was then added at a concentration of 5 µg/mL to every well on the plate, and pseudovirus, diluted to a titer of 1 x 10^8^ total integrated intensity units/mL as per titers determined independently for each pseudovirus batch, was added 1:1 to the diluted serum samples. The plates were incubated for 1 hour at 37°C and 5% CO_2_. Serum-virus mix was then added in duplicate to Human Angiotensin-Converting Enzyme 2 (hACE2)-expressing HEK-293 cells (BEI Resources, #NR-52511) seeded at 4 x 10^5^ cells/mL on a 96-well flat-bottom plate and incubated at 37°C and 5% CO_2_ for 72 hours. To determine 50% inhibitory concentration (IC_50_) values, plates were scanned on a high-content fluorescent imager (ImageXpress Pico Automated Cell Imaging System, Molecular Devices, San Jose, CA) for ZsGreen expression. Total integrated intensity per well was used to calculate the percent of pseudovirus inhibition in each well. Neutralization data for each sample were fit with a four-parameter sigmoidal curve that was used to interpolate IC_50_ values.

A Wuhan-strain pseudoneutralization test using the World Health Organization (WHO) standard for neutralization assays with an official IC_50_ of 1000 resulted in an IC_50_ of 7800 using our assay, suggesting that our test is more sensitive than the WHO assay but may overreport IC_50_ values—by less than one log_10_—an effect likely most pronounced for strongly neutralizing samples.

### Splenocyte harvest, intracellular cytokine staining, and flow cytometry

Spleens were dissociated in 4 mL of RPMI medium by manual maceration through a cell strainer using the end of a syringe plunger. Homogenized samples were centrifuged at 400 x g for 5 minutes to pellet red blood cells. The supernatants containing lymphocytes were transferred to 5-mL mesh-cap tubes to strain out any remaining tissue debris or were lysed with ammonium-chloride-potassium (ACK) buffer and washed. Cell counts for each sample were obtained on a Guava easyCyte (Luminex, Austin, TX). Each spleen sample was seeded in 96-well round-bottom plates at 1-2 x 10^6^ cells per well in RPMI medium containing 10% FBS, 50μM beta-mercaptoethanol, CD28 costimulatory antibody, and brefeldin A. Cells were stimulated with one of three stimulation treatments: 0.0475% dimethyl sulphoxide (DMSO) as a negative stimulation control, 10 μg/well (1 μg/mL) of spike peptide pool (JPT Peptide Technologies, Berlin, Germany; #PM-WCPV-S-1) in an equivalent amount of DMSO, or 10 μg/well of phorbol myristate acetate (PMA)/ionomycin solution. After 6 hours of incubation at 37°C with 5% CO_2_, plates were centrifuged at 400 x g for 3 minutes, the supernatants were removed by pipetting, and cells were resuspended in PBS. Plates were centrifuged, the supernatants were removed, and cells were stained for flow cytometry. Splenocytes were stained for viability with Zombie Green (BioLegend, San Diego, CA) in 50 μL of PBS, and then Fc receptors were blocked with CD16/CD32 antibody (Invitrogen). Cells were then surface stained with fluorochrome-labeled mAbs specific for mouse CD4, CD8, CD44, and CD107a in 50 μL of staining buffer (PBS with 0.5% bovine serum albumin and 0.1% sodium azide). Cells were washed twice, permeabilized using the Fixation/Permeabilization Kit (BD Biosciences, Franklin Lakes, NJ), and stained with fluorochrome-labeled mAbs specific for mouse TNFα, IL-2, IFNγ, IL-5, IL-10, and IL-17A. After two washes in staining buffer, cells were resuspended in 100 μL of staining buffer and analyzed on an LSRFortessa flow cytometer (BD Biosciences). After initial gating for live CD4^+^ or CD8^+^ lymphocytes, cells were gated for cytokine positivity. Quality of the response was determined by gating on cells that were double or triple positive for these markers. Cells triple positive for TNFα, IL-2, and IFNγ were considered activated polyfunctional T cells.

### T-cell ELISpot assay

ELISpot plates (MilliporeSigma) were coated with either IFNγ (BD Biosciences, #51-2525KZ), IL-17A (Invitrogen, #88-7371-88), or IL-5 (BD Biosciences, #51-1805KZ) capture antibodies at a 1:200 dilution in ELISpot coating buffer (eBioscience, Waltham, MA). After an overnight incubation at 4°C, plates were washed and then blocked with complete RPMI (cRPMI) medium for at least 2 hours. Splenocytes harvested as described above were plated at 2 x 10^5^ cells per well. A subset of each sample was stimulated with PepMix SARS-CoV-2 (JPT Peptide Technologies, #PM-WCPV-S-1) at a final concentration of 1 µg/mL. Plates were then incubated at 37°C and 5% CO2 for 48 hours. After a wash with PBS with 0.1% Tween 20, 100 µL of detection antibody (IFNγ, BD Biosciences, #51-1818KA; IL-17A, Invitrogen, #88-7371-88; and IL-5, BD Biosciences, #51-1806KZ) was added at a 1:250 dilution in ELISpot diluent (eBioscience) overnight at 4°C. Plates were washed and developed with Vector NovaRED Substrate Peroxidase (Vector Laboratories, Burlingame, CA; #SK-4800) for 15 minutes. The reaction was stopped by washing the plates with deionized water, and plates were left to dry in the dark. Spots were counted and data were analyzed using ImmunoSpot software (Cellular Technology Limited, Cleveland, OH).

### Bone marrow IgA and IgG antibody-secreting cell ELISpot assays

Presence of antibody-secreting bone marrow-resident cells were measured by ELISpot. MultiScreen IP Filter plates (0.45 µm, MilliporeSigma) were treated with 15 µL of 35% ethanol for 30 seconds. After a wash with PBS, plates were coated with 100 µL of Recombinant SARS-CoV-2 Spike His Protein, Carrier Free, (R&D Systems, #10549-CV-100) at a concentration of 2 µg/mL diluted in ELISpot coating buffer (eBioScience). Plates were incubated overnight at 4°C, washed with PBS with 0.1% Tween 20, and blocked with cRPMI medium for at least 2 hours.

To prepare bone marrow, both femurs were removed from each mouse and inserted into a snipped-end 0.6 mL tube (Eppendorf, Hamburg, Germany) inserted into a 1.5 mL Eppendorf tube containing 1 mL of cRPMI medium. Femurs were centrifuged for 15 seconds at 10,000 rpm, and supernatant was discarded. The cell pellets were briefly vortexed and resuspended in 200 µg of RBC lysis buffer (Invitrogen), and they were incubated on ice for 30 seconds. After addition of 800 μL of cRPMI medium, cells were centrifuged 5 minutes at 400 x g, and supernatant was decanted. Cells were resuspended in 1 mL of cRPMI medium, counted, and transferred to prepared filter plates described above at 1 million cells per well followed by a 3-fold dilution across five adjacent wells.

After a 3-hour incubation, plates were washed with PBS with 0.1% Tween 20, and secondary antibody (Goat Anti-Mouse IgG-HRP or IgA-HRP [SouthernBiotech, Birmingham, AL; #1030-05 and #1040-05]) was added at a 1:1000 dilution in PBS with 0.1% Tween and 5% FBS overnight at 4°C. Plates were then washed three times in PBS with 0.1% Tween 20 and two times in PBS. For development, 100 µL of Vector NovaRED Substrate Peroxidase (Vector Laboratories, #SK-4800) was applied for 7 minutes. The reaction was stopped by rinsing plates with distilled water for 2 minutes, and plates were dried in the dark. Spots were counted and data were analyzed using ImmunoSpot software (Cellular Technology Limited).

### Statistical analyses

Log-normalized pseudovirus variant neutralization was compared using mixed effects analysis with multiple comparison correction. SARS-CoV-2-specific IgG levels in mouse sera by stored vaccines were analyzed on log-transformed data by one-way ANOVA at each timepoint followed by Dunnett’s multiple comparison test comparing each stored sample to the freshly complexed positive control at each timepoint. All statistical analyses were conducted using Prism 9 (GraphPad Software, San Diego, CA).

## ACKNOWLEDGMENTS

We would like to thank Jesse Bloom (Fred Hutchinson Cancer Research Center) and Thomas Peacock (Imperial College London) for sharing the SARS-CoV-2 spike protein plasmids used for pseudovirus production. We are additionally grateful to Elise Larson, who conducted the manufacture of the NLC used in these studies; Richard Cabullos, who manufactured some RNA products used in this study; and Jeff Guderian. John Tite provided valuable technical input. We also thank Theresa Britschgi, who was instrumental as project manager for this work; Valerie Soza, for editorial assistance; and Cassandra Baden, for graphic design.

This work was funded in whole or in part with funds from Amyris, Inc., and ImmunityBio, Inc. The content is solely the responsibility of the authors and does not necessarily represent the views of the funders.

## AUTHOR CONTRIBUTIONS

E.A.V., A.G, D.H., C.J.P., C.B.F., and C.C conceived the study. E.A.V., A.G., C.J.P., C.B.F., and C.C. supervised the research and administered the project. E.A.V., C.B.F., and C.C. acquired funding. C.J.P. and P.S.S. provided funding. E.A.V., A.G., D.H., M.F.J., P.B., S.R., J.S., R.M., J.B., S.B., C.P., and J.G. contributed to the investigation. E.A.V., A.G., D.H., M.F.J., P.B., S.R., R.M., and S.B. contributed to methodology. D.H., M.F.J., P.B., and S.B. curated the data. E.A.V., A.G., D.H., M.F.J., P.B., S.R., J.S., and S.B. analyzed the data. E.A.V., A.G., D.H., M.F.J., and J.S. visualized the data. E.A.V., A.G., M.F.J., and C.C. wrote the first draft of the manuscript. All authors reviewed and edited the manuscript.

## DECLARATION OF INTERESTS

C.B.F. is co-inventor on patent applications relating to PCT/US2018/37783, “Nanostructured Lipid Carriers and stable emulsions and uses thereof.” A.G. and E.A.V. are co-inventors on U.S. patent application nos. PCT/US21/40388, “Co-lyophilized RNA and Nanostructured Lipid Carrier,” and 63/144,169, “A thermostable, flexible RNA vaccine delivery platform for pandemic response.” C.J.P. owns shares and possesses stock options in Amyris, Inc. P.S.S. owns shares of ImmunityBio, Inc. All other authors declare they have no competing interests.

