## Supplemental Information for "A self-amplifying RNA vaccine against COVID-19 with long-term room-temperature stability"

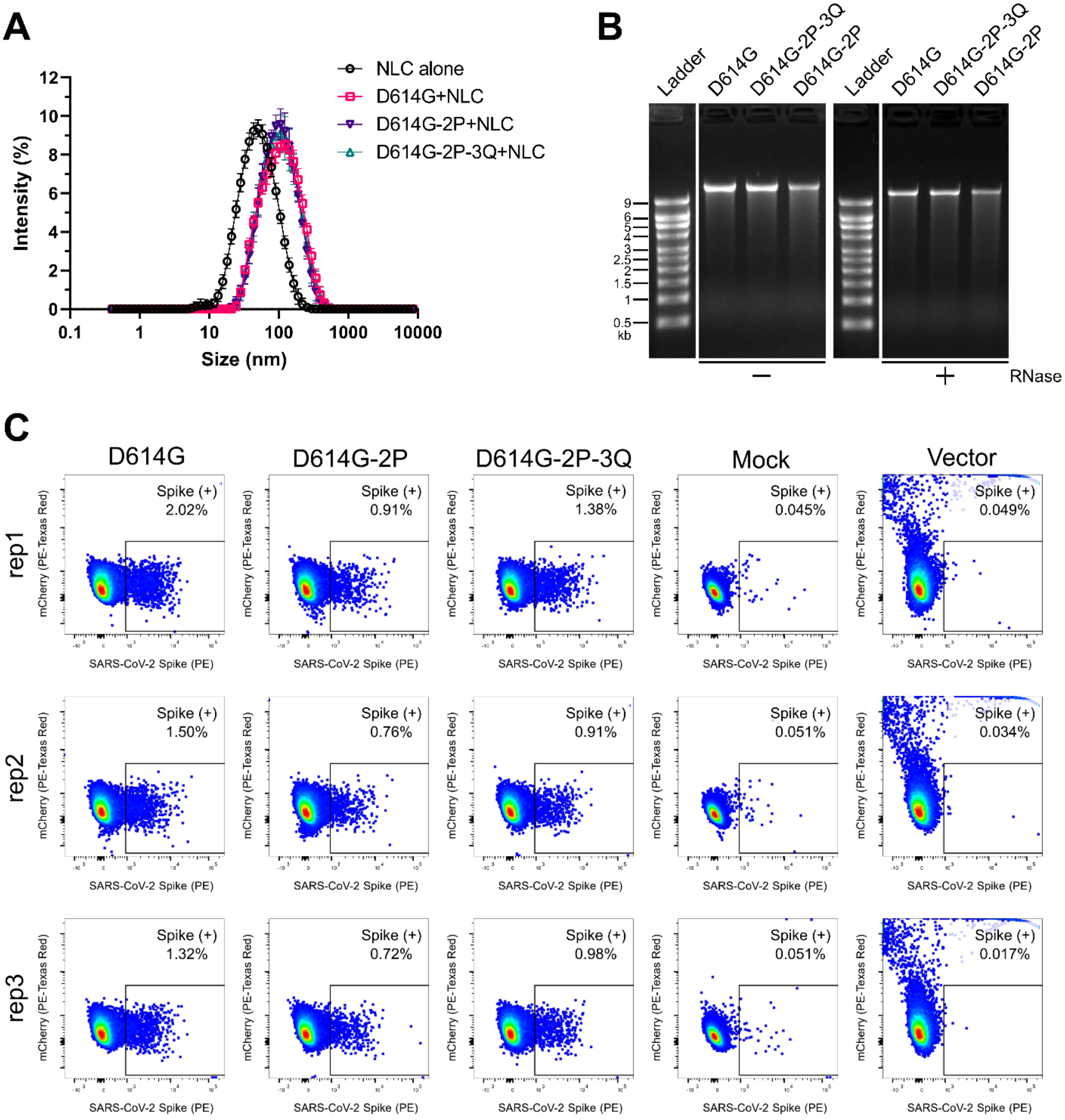


**Supplemental Figure S1. Characterization of SARS-CoV-2 saRNA/NLC vaccine constructs D614G (baseline), D614G-2P, and D614G-2P-3Q (AAHI-SC2).** A) Size distribution of NLC particles alone and saRNA/NLC vaccine particles after complexing with each saRNA construct. B) Each saRNA construct complexed to NLC particles is protected from RNase degradation. C) Flow cytometric detection of SARS-CoV-2 spike expression by HEK-293 T cells after *in vitro* transfection of 500 µg saRNA/NLC per well of a cell-seeded 12-well plate. Cells were harvested 24 hours post-transfection and stained with a Cy3-conjugated rabbit polyclonal antibody to SARS-CoV-2 spike (Abcam, Cambridge, UK; #AB272504). Flow cytometry data for the three saRNA/NLC vaccine constructs and two controls (mock and mCherry reporter vector) include three biological transfection replicates. Cells transfected with the mCherry reporter vector/NLC demonstrate expression of mCherry but no background SARS-CoV-2 staining as expected.

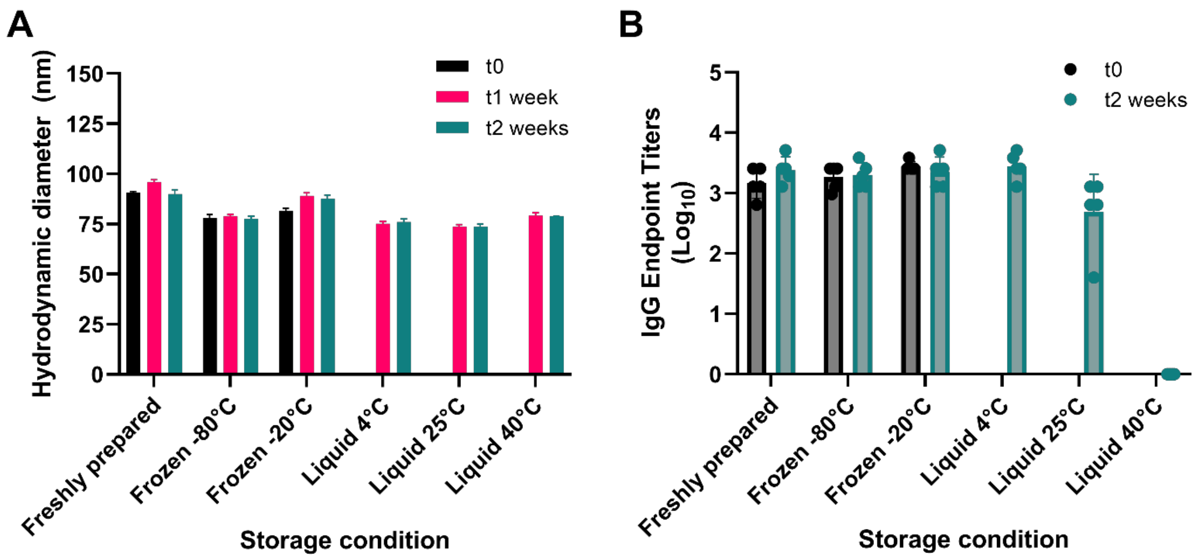


**Supplemental Figure S2. Stability of the liquid SARS-CoV-2 saRNA/NLC vaccine (AAHI-SC2) after 2 weeks of storage at different temperatures.** A) Hydrodynamic (Z-average) diameter of the vaccine complex prior to or after storage at the given conditions. B) Serum SARS-CoV-2 spike protein-binding IgG induced in C57BL/6 by a single 10µg dose of vaccine after storage at the indicated conditions. *n* = 5 mice/group. Serum samples were taken 2 weeks after vaccine injection and analyzed by ELISA.

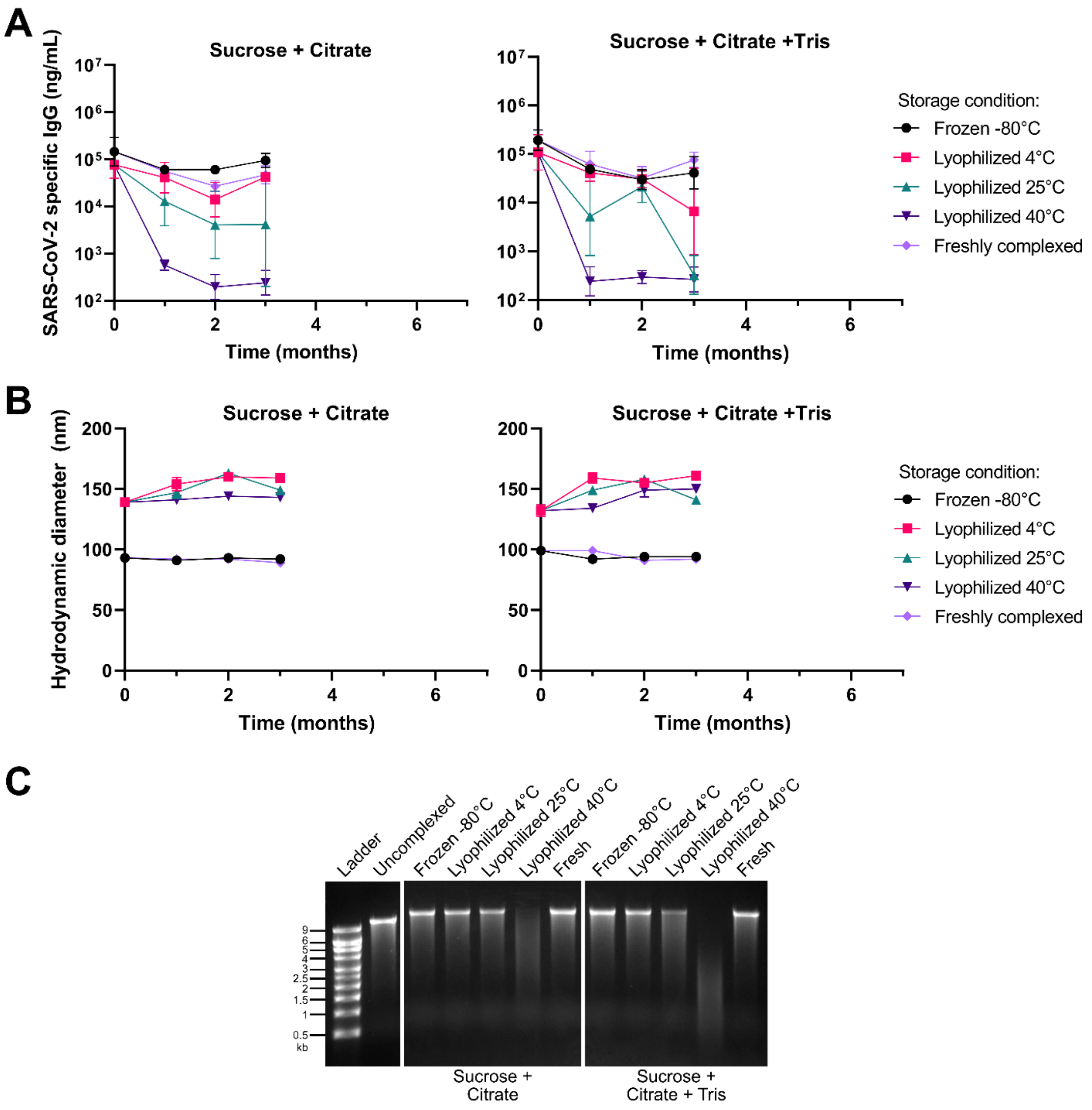


**Supplemental Figure S3. Stability of the lyophilized SARS-CoV-2 saRNA/NLC vaccine (AAHI-SC2) in alternative excipient backgrounds after 3 months of storage at different temperatures.** A) Serum SARS-CoV-2 spike protein-binding IgG after vaccine storage under the indicated conditions. *n* = 5 mice/group. B) Hydrodynamic (Z-average) diameter of the vaccine complex. C) Integrity of vaccine RNA after 3 months of storage under the indicated conditions. Sucrose + citrate solution comprised 20% w/v sucrose and 5mM sodium citrate; sucrose + citrate + Tris comprised 20% sucrose, 1mM sodium citrate, and 10mM Tris.

**Supplementary Table S1. Physicochemical stability of NLCs using DOTAP from different vendors, stored at 2-8°C.**

| | Time point | Particle diameter (Z-ave, nm) | Size polydispersity index (PDI) | Visual appearance | pH | DOTAP content (mg/mL) | Squalene content (mg/mL) |
|---|---|---|---|---|---|---|---|
| **Expected range** | -- | 40 +/- 20 | for information only | white, translucent, single phase | 5.0-6.5 | 30.0 +/- 9.0 | 37.5 +/- 7.5 |
| **DOTAP vendor 1** | T = 0 | 38.3 +/- 0.8 | 0.19 +/- 0.01 | white, translucent, single phase | 5.75 | 25.1 +/- 1.0 | 32.9 +/- 0.3 |
| | T = 3 mon | 42.0 +/- 0.3 | 0.26 +/- 0.02 | white, translucent, single phase | 5.63 | 25.8 +/- 0.3 | 33.6 +/- 0.3 |
| **DOTAP vendor 2** | T = 0 | 46.9 +/- 1.1 | 0.28 +/- 0.00 | white, translucent, single phase | 5.67 | 24.2 +/- 1.8 | 31.0 +/- 1.4 |
| | T = 3 mon | 48.8 +/- 1.0 | 0.28 +/- 0.01 | white, translucent, single phase | 5.63 | 26.8 +/- 0.9 | 35.1 +/- 0.6 |
